# CUDASW++4.0: Ultra-fast GPU-based Smith-Waterman Protein Sequence Database Search

**DOI:** 10.1101/2023.10.09.561526

**Authors:** Bertil Schmidt, Felix Kallenborn, Alejandro Chacon, Christian Hundt

## Abstract

**Background:** The maximal sensitivity for local pairwise alignment makes the Smith-Waterman algorithm a popular choice for protein sequence database search. However, its quadratic time complexity makes it compute-intensive. Unfortunately, current state-of-the-art software tools are not able to leverage the massively parallel processing capabilities of modern GPUs with close-to-peak performance. This motivates the need for more efficient implementations.

**Results:** CUDASW++4.0 is a fast software tool for scanning protein sequence databases with the Smith-Waterman algorithm on CUDA-enabled GPUs. Our approach achieves high efficiency for dynamic programming-based alignment computation by minimizing memory accesses and instructions. We provide both efficient matrix tiling, and sequence database partitioning schemes, and exploit next generation floating point arithmetic and novel DPX instructions. This leads to close-to-peak performance on modern GPU generations (Ampere, Ada, Hopper) with throughput rates of up to 1.94 TCUPS, 5.01 TCUPS, 5.71 TCUPS on an A100, L40S, and H100, respectively. Evaluation on the Swiss-Prot, UniRef50, and TrEMBL databases shows that CUDASW++4.0 gains over an order-of-magnitude performance improvements over previous GPU-based approaches (CUDASW++3.0, ADEPT). In addition, our algorithm demonstrates significant speedups over top-performing CPU-based tools (BLASTP, SWIPE, SWIMM2.0), can exploit multi-GPU nodes with linear scaling, and features an impressive energy efficiency of up to 15.7 GCUPS/Watt.

**Conclusion:** CUDASW++4.0 changes the standing of GPUs in protein sequence database search with Smith-Waterman alignment by providing close-to-peak performance on modern GPUs. It is freely available at https://github.com/asbschmidt/CUDASW4.

## Background

Computing optimal local pairwise alignments of biological sequences with the dynamic programming (DP)-based Smith-Waterman (SW) algorithm [1] is a core algorithm in bioinformatics. A prominent example is protein sequence database search where similarities between a query sequence and a database sequence can be identified by computing their optimal local alignment score. However, the associated time complexity to compute an alignment score between a pair of sequences is proportional to the product of their lengths which renders Smith-Waterman a comparably expensive yet highly accurate algorithm. The computational demand is further increased when considering the ever-growing number of sequences stored in largescale datasets propelled by the continuous advances in high-throughput technologies.

Consequently, a number of parallelized implementations have been developed for computing pairwise sequence alignments on a variety of architectures including CPUs [2, 3, 4, 5, 6], GPUs [7, 8, 9, 10, 11, 12, 13, 14], and FPGAs [15, 16, 17, 18]. They typically target two types of application scenarios:

1. computing a single pairwise alignment of two very long DNA sequences (typically whole genomes or chromosomes), and
2. computing batches of different pairwise alignments of DNA/RNA or protein sequences of shorter length.

Widely adopted parallelization schemes are *inter-sequence* (computes a number of independent alignment tasks in parallel), *intra-sequence* (computes the DP matrix of a single alignment in parallel), and their *hybrid* combination.

Another approach to accelerate time-consuming sequence database search is the usage of fast heuristics to replace expensive alignments. Prominent examples include FASTA [19] and BLAST [20, 21, 22]. These heuristics can often produce reasonably good results, but could fail to detect some distantly related sequences due to the loss of sensitivity [23, 24, 25]. In addition, their efficient fine-grained parallelization is made challenging by irregular computation and memory access patterns for vector and streaming processing architectures.

Existing GPU implementations specifically optimized for protein sequence database search include CUDASW++3.0 [8], ADEPT [12], SWhybrid [14], and NVBIO [13]. They typically compute an independent alignment per CUDA thread block whereby threads compute cells along a minor diagonal of the DP matrix in parallel. Data between neighboring diagonals is typically communicated via global or shared memory. Unfortunately, they are all limited by inefficient memory access schemes and thus cannot fully exploit the performance of modern GPUs leading to slow runtimes.

In this paper, we present CUDASW++4.0, which yields significantly faster and highly energy-efficient SW-based protein database searches compared to previous GPU implementations. To achieve high performance on modern GPUs, we take advantage of the following concepts:

- fine-grained parallelization strategy based on warp intrinsics for fast inter-thread communication (instead of using shared or global memory) and minimizing overall executed instructions,
- tiling scheme for DP matrices in order to support batches with highly varying sequence lengths occurring in typical real-world database,
- sequence database partitioning to achieve fully-functional and high-performance support on GPUs with limited memory size,
- advanced sequence length binning strategies combined with warp specialized kernels,
- efficient support of new architectural features of modern GPUs, such as half-precision using half2 arithmetic or s16x2 arithmetic by using novel DPX instructions.

We initially evaluate the theoretical peak performance of CUDASW++4.0 using simulated databases comprised of equal-length protein sequences for different datatypes (float, half2, int32, s16x2) on three different recent GPU architectures (Ampere, Ada, Hopper). This also includes an evaluation of the performance of the recently introduced DPX instructions [26]. Subsequently, we evaluate the performance of CUDASW++4.0 using the real-world Swiss-Prot, UniRef50, and TrEMBL databases. Our performance comparison shows that CUDASW++4.0 consistently outperforms other GPU-based implementations (CUDASW++3.0, ADEPT) and CPU-based tools (SWIPE, SWIMM2.0, BLASTP) for queries of various lengths. CUDASW++4.0 achieves peak throughput rates of up to 1.94 TCUPS (<u>T</u>rillions of <u>C</u>ell <u>U</u>pdates <u>P</u>er <u>S</u>econd), 5.01 TCUPS, and 5.71 TCUPS on an A100, L40S, and H100 GPU, respectively. In addition, even higher speeds can be achieved by using multiple GPUs connected to the same host with close-to-linear linear scaling; e.g. 96.5% parallel efficiency can be achieved for a server with 8 H100 GPUs. Furthermore, our conducted power consumption analysis shows that GPUs can offer an order-of-magnitude higher energy efficiency than the most recent FPGA implementation at the same power budget.

### Smith-Waterman Algorithm

Consider two sequences *Q* = (*q*_0_*q*_1_ … *q*_*m*−1_) and *S* = (*s*_0_*s*_1_ … *s*_*n_*1_) over the alphabet Σ of length *m* and *n*, respectively. For each pair (*q*_*i*_, *s*_*j*_) of characters the recurrence relation in Eq. 1 of the DP algorithm for local alignment decides if these characters should be aligned, if a gap should be inserted, or a new alignment is started. A corresponding score is recorded in DP matrix *H*.

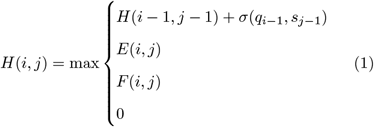

where 1 ≤ *i* ≤ *m*, 1 ≤ *j* ≤ *n* and

- *E* and *F* are defined in Eqs. 2 and 3 and determine the *gap penalty scheme*,
- *σ* is a substitution function over Σ × Σ that determines the score of aligning two characters (for protein sequences this is typically a BLOSUM or PAM matrix).

We consider an *affine gap penalty* scheme, where a gap of length *k* is penalized by *α* + (*k* − 1) · *β* where *α* is the cost of the first gap (opening) and *β* the cost of each subsequent gap (extension):

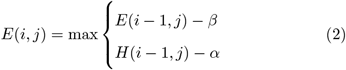

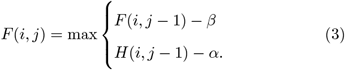

The matrices are initialized by *H*(0, 0) = *H*(*i*, 0) = *H*(0, *j*) = 0, *E*(0, 0) = *F* (0, 0) = *F* (*i*, 0) = *E*(0, *j*) = −∞, *E*(*i*, 0) = −*α* − (*i* − 1) · *β*, and *F* (0, *j*) = −*α* − (*j* − 1) · *β*.

The dependency relation in *H* is illustrated in Fig. 1, where light gray cells are initialization cells and dark gray cells indicate the ancestral subproblems of the currently active cell in black.

**Figure 1.**
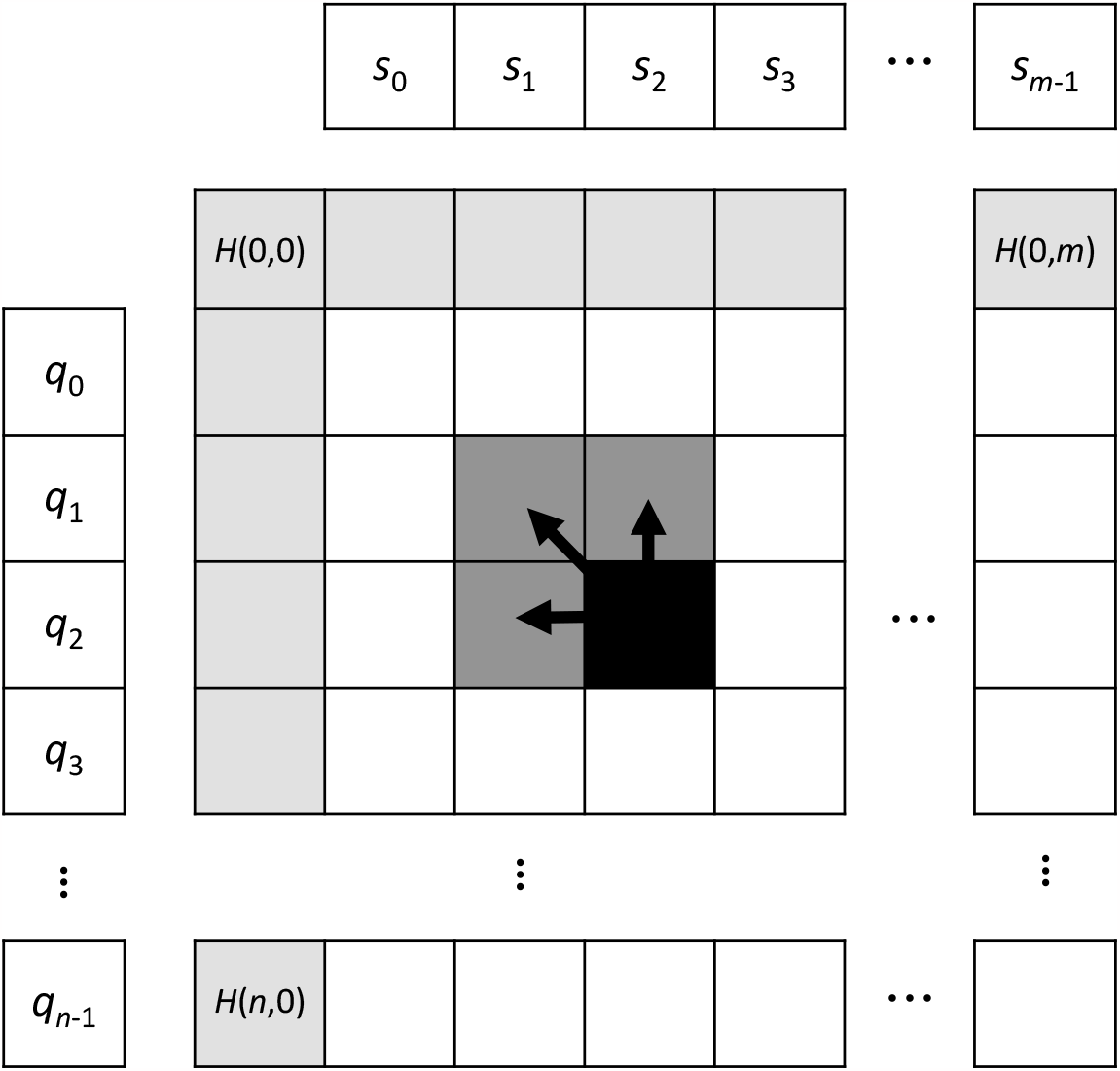
Layout and dependency relationships of the DP matrix *H*.

The optimal local alignment score is equal to the maximum value in *H* and can be computed in linear space 𝒪(min{*m, n*}) and quadratic time 𝒪(*m*·*n*). Note that for protein database search we are interested in score-only computation but not the traceback.

### Features of CUDA-enabled GPUs

CUDA kernels are executed using a number of independent thread blocks. Each thread block is mapped onto exactly one streaming multiprocessor (SM) and consists of a number of warps. All 32 threads within a warp are executed in lock-step fashion (SIMT).

CUDA-enabled GPUs contain several types of memory: large but high latency global memory and fast but small on-chip shared and constant memory. Nevertheless, the fastest way to access data is through usage of the thread-local register file.

Modern GPUs provide instructions for warp-level collectives in order to efficiently support communication of data stored in registers between threads within a warp without the need for accessing global or shared memory.

A crucial feature of our approach is the usage of warp shuffles for low latency communication and minimization of memory traffic. In particular, we take advantage of the warp-level collectives __`shfl_down_sync()` and __`shfl_up_sync()`; e.g. the intra-warp communication operation `R1 = __shfl_up_sync(0xFFFFFFFF, R0, 1, 32);` moves the contents of register `R0` in thread *i* within each warp to register `R1` in thread *i* + 1 for 0 ≤ *i <* 31.

### Advanced Architectural GPU features for DP

The three latest GPU generations (Ampere, Ada, Hopper) have incrementally included key hardware features that can be used to improve the performance of DP algorithms. Each architecture includes the following hardware forward-compatible features (e.g. Hopper supports all points below):

1. `[Ampere] Half2:` Support of half-precision floating point arithmetic instructions that allow for the simultaneous execution of one 32-bit operation on two half-precision values. This allows for computing two values of the DP recurrence relation within a single register, effectively doubling the throughput. Ampere enables the necessary `half2 min/max` comparison function present in SW.
2. `[Ada] FP dual-port and max3:` Enables dual ports for min/max and add/sub floating point (FP) operations. This allows us to execute both instructions independently in the SM and increase the overall throughput. Additionally, Ada introduced a new half2 set of instructions to perform the min/max operation of three input values (`vhmnmx`).
3. `[Hopper] ALU dual-port and DPX:` Inclusion of DPX instructions to support fast computation of typical DP recurrence relations with integer-based arithmetic. This includes a set of functions that enable finding a max value of three inputs (`vimax3`), as well as fused addition with max operations (`viaddmax`). The instructions support any combination of min/max, sub/add for 16-bit or 32-bit integer parameters, with optional ReLU (clamping to zero). Similarly to Ada, Hopper supports integer parallel execution of max/min and add/sub in independent SM internal resources.

Following sections will describe the necessary algorithmic changes to achieve high performance using these features and analyse their impact on real workloads.

## Implementation

### Program Information and Workflow

CUDASW++4.0 is written in C++17 and targets Linux workstations and servers. It gains high speed by benefiting from the use of GPU accelerators attached to a host CPU (typically via PCIe) and requires a system with one or more GPUs compatible with CUDA 12. Detailed instructions are included in the software repository.

Our algorithm for scanning a protein sequence database (DB) with a (single) query protein sequence *Q* on a single GPU consists of the following stages:

1. Read local DB and *Q* from file.
2. Transfer the first batch of DB sequences and *Q* from CPU to GPU.
3. Partition current DB batch by sequence lengths on GPU.
4. Execute GPU alignment kernels for each partition.
5. Transfer the next batch of DB sequence from CPU to GPU concurrently to alignment computation.
6. Re-compute all alignments with indicated overflow (for 16-bit arithmetic) using GPU alignment kernels with 32-bit arithmetic.
7. If all DB sequences have been aligned to *Q* continue with Stage 8, otherwise continue with Stage 3.
8. Sort all alignment scores on GPU and transfer results to CPU for output.

Since GPU memory can be a scarce resource, we process batches of DB sequences in order to support large-scale protein sequence sets. Furthermore, this approach allows for concurrent PCIe data transfer and kernel execution.

Although libraries for parsing of FASTA files exist [27], this task can still take significant time for reading databases like UniRef or TrEMBL. Thus, CUDASW++4.0 includes a preprocessor called `makeDB` (Stage 0) that creates a local database from protein FASTA files. We also support reading input files in gzip-compressed format (fasta.gz). The local database can then be used by CUDASW++4.0 for scanning with greatly accelerated file parsing time^[1]^. This approach is effective since databases are frequently scanned but only need to be preprocessed once. Sequences in the database are stored in a custom format and are sorted by length to allow for partitioning by sequence length during the main algorithm.

Note that our program also accepts a set of query sequences {*Q*_0_, … *Q*_*P* −1_} in a single FASTA file as input. In the case of multiple query sequences the workflow is repeated (except for reading the input files) for each query. If all database sequences can be stored in GPU device memory (which is typically the case for relatively small or medium-sized databases like Swiss-Prot or UniRef50) they only need to be transferred once for all *P* scan tasks. Figure 2 illustrates the workflow of CUDASW++4.0.

**Figure 2.**
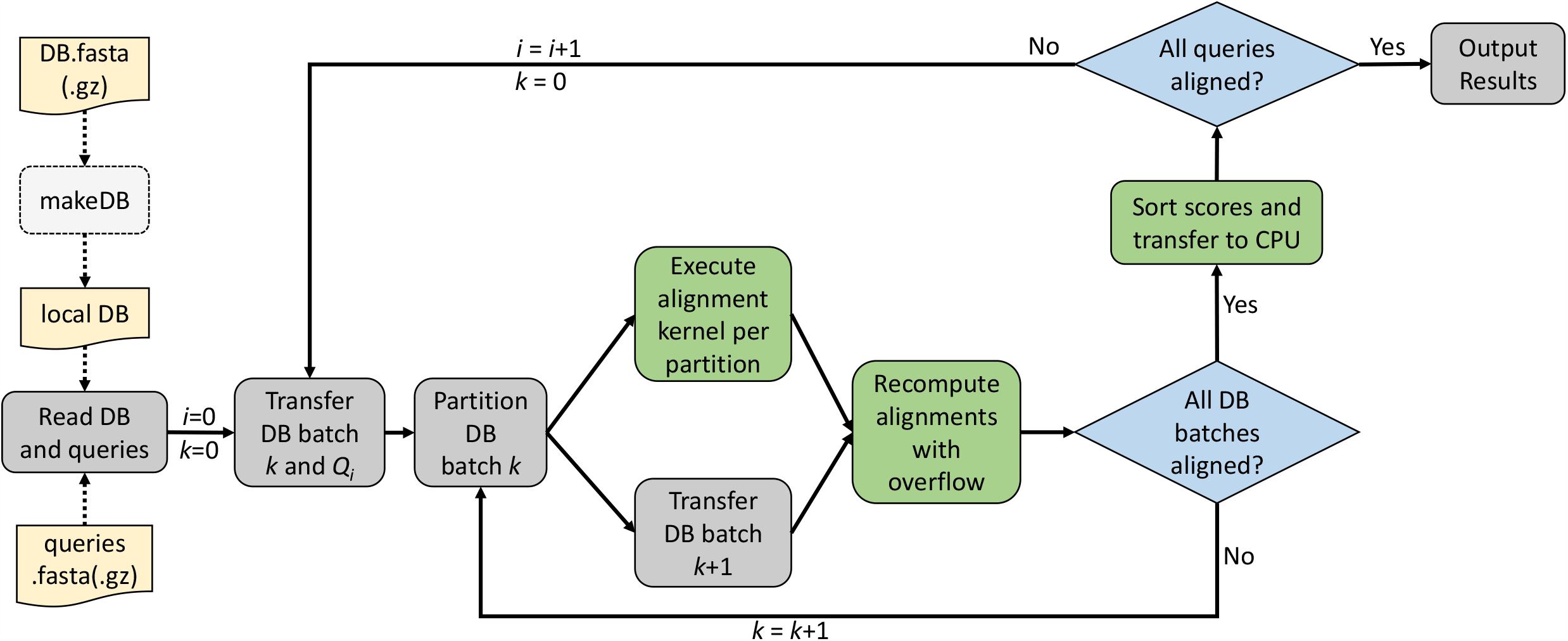
Workflow of CUDASW++4.0 for scanning a database DB with a set of queries *{Q*_0_, … *Q*_*P* −1_*}*. Steps executed on the GPU (CPU) are colored in green (grey).

In addition to the features shown in the standard workflow we offer an *interactive* mode and a *multi-GPU* mode. In interactive mode the user can supply query sequences from the command line to scan a (pre-loaded) database. The multi-GPU mode takes advantage of several GPUs attached to the same host CPU by partitioning the input database into equal-sized parts per GPU.

### GPU-based Alignment Computation

We base our parallelization scheme on computing an independent alignment per (sub)warp – a group of synchronized threads executed in lockstep that can efficiently communicate by means of warp shuffles. Threads in a (sub)warp compute DP matrix cell values in cooperative fashion. Note that according to Eqs. 1, 2, 3 each DP cell depends on its left, upper, and upper-left neighbour (see Fig. 1), which means that (parallel) computation of DP cells has to follow this topological order.

The fine-grained parallel design proposed in this work combined with optimizations described in the following sections leads to important advantages [28, 29] such as (a) avoiding compute- and memory-thread divergence, (b) dynamic partitioning of large memory footprints on local memories, and (c) reduction of the overall executed instructions (e.g. removing LD/ST instructions) thus allowing the implementation of highly efficient tiling methods.

In order to unlock the full potential of modern GPUs, we apply the following techniques:

- Full in-register computation of recurrence relations.
- Low-latency communication of neighboring DP cells between threads through warp shuffles.
- Reduction of frequent sequence character loading from memory by employing an intra-warp communication scheme based on warp shuffles.
- Efficient scheme for substitution table lookups *Σ*(*q*_*i*_, *s*_*j*_) for BLOSUM matrices.
- DP matrix tiling scheme for long sequences.
- Optimized kernel variants for various sequence lengths and datatypes using half2 and float arithmetic and evaluation of DPX instructions for integer arithmetic (int32 and s16x2).

#### Mapping DP Matrix Cells to Threads

Each (sub)warp (a group of *p* threads *T*_0_, …, *T*_*p*−1_ with *p* ∈ {2, 4, 8, 16, 32} executed in lock step), computes the optimal local alignment score between a (query) protein sequence *Q* and a database sequence *S*^(*i*)^. Assume *Q* = (*q*_0_*q*_1_ … *q*_*m*−1_) of length *m* and 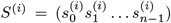 of length *n* = *k* · *p*. We assign *k* columns of the DP matrix to be calculated by each thread. Computation proceeds along a *wavefront* in *m* + *p* iterations. In iteration *i*, thread *T*_*t*_ computes *k* adjacent cells of the DP matrix row *i* − *t*. In practice, the maximum possible value of *k* depends on the number of available registers and used datatype. Each DP matrix cell depends on its left, upper, and upper-left neighbor. All cells of the current and previous iteration are stored in thread-local registers. The required access of thread *T*_*t*_ to the rightmost value of thread *T*_*t*−1_ computed in the previous iteration is accomplished by using the low-latency warp shuffle instruction __`shfl_up_sync()`. Our mapping strategy is illustrated in Fig. 3.

**Figure 3.**
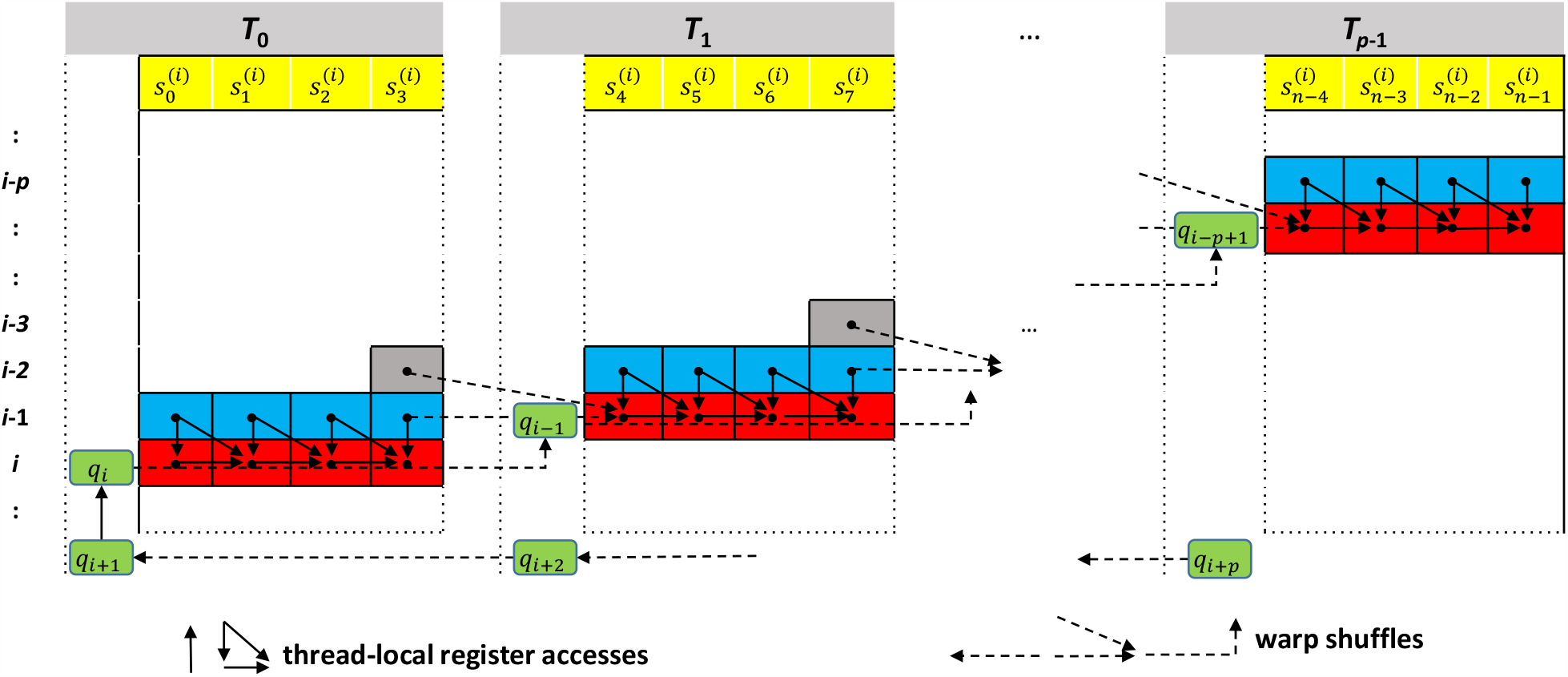
Example of mapping DP matrix computation for comparing *Q* = (*q*_0_*q*_1_ … *q*_*m*−1_) length *m* and 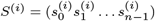 of length *n* = *k · p* to a group of *p* threads using *k* = 4. Each thread computes *k* matrix cells (red) in each iteration *i* using the values of the antecedent row (blue) and an additional value from the second antecedent row (gray). Threads communicate values in the rightmost of their *k* columns as well as characters of *q* (green) by warp shuffle operations (dashed arrows).

At the beginning each thread *T*_*t*_, 0 ≤ *t < p* loads *k* characters of *S*^(*i*)^ (i.e.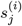, *t* · *k* ≤ *j < t* · (*k* + 1)) from global device memory. Note that the loaded characters of *S*^(*i*)^ remain the same for each column of the DP matrix while the required characters from *Q* vary. In order to avoid repeated, expensive reading from global device memory, we store *Q* in CUDA constant memory and only load new characters from *Q* into registers every *p* iterations. They are exchanged between threads via warp shuffles by means of using two registers *c*_*q*_*_*_current_ and *c*_*q*_*_*_next_. Register *c*_*q*_*_*_current_ always stores the query character needed for the current computation by each thread. At the start of each iteration, thread 0 copies the current value of *c*_*q*_*_*_next_ to *c*_*q*_*_*_current_. Afterwards, a warp shuffle-up and a warp shuffle-down update *c*_*q*_*_*_current_ and *c*_*q*_*_*_next_, respectively, with characters from neighboring threads. Only in iterations 0 ≤ *i < m* + *p* with *i* mod *p* = 0 each Thread *t* loads a new query character *q*_*i*+*t*_ from memory and stores it in *c*_*q*_*_*_next_. Note that by reading `char4` values we are able to further reduce the frequency of loading values of *Q* by a factor 4.

CUDASW++3.0 [8] created a sequence-specific *scoring profile* that is used for performing substitution table lookups 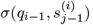. However, the associated space complexity of O(|Σ| · *n*) per warp makes it unattractive for large values of *n*. Thus, *Σ* of size O(|Σ| · |Σ|) is simply loaded into shared memory by all threads of a CUDA thread block in cooperative fashion instead. The two-dimensional lookup is then efficiently performed in two stages:

1. A row is selected by the current character of the query sequence stored; i.e. *r* = *Σ*(*c*_*q*_*_*_current_, −).
2. This row is then looked up in *k* subsequent iterations using the *k* letters of the database sequence in each thread; i.e.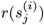, *t* · *k* ≤ *j < t* · (*k* + 1)).

For half precision arithmetic, the substitution table is stored as a matrix of size |Σ| · |Σ|^2^, which allows for efficient lookup of two 16-bit values within a 32-bit word simultaneously and still fits into shared memory. Note that all warps within a CUDA thread block access the same substitution table and thus do not require additional shared memory resources (as would the case for the *scoring profile* used in CUDASW++3.0).

To compute *k* DP matrix cells, each thread first looks up the substitution scores and subsequently computes the recurrence relations (Eq. 1–3). This involves several maximum, addition, and subtraction operations using only thread-local in-register computation. Subsequently, the rightmost DP cell computed in Thread *t* is communicated to Thread *t* + 1 using the low latency warp shuffle instruction __`shfl_up_sync()`. Neighboring query sequence letters are also communicated between threads in the same way (see Fig. 3).

Each thread continuously tracks the maximum value of its computed values of *H* in a register. At the end of each pairwise alignment computation the overall maximum score is determined by a warp-level maximum-reduction operation, which is subsequently written to CUDA global memory as output.

#### DP Matrix Tiling Scheme

The maximum number of registers per thread is typically limited to 255 on CUDA-enabled GPUs. This restricts the maximum number of columns (*k*_*max*_) that can be efficiently computed by one thread. For our implementation, we observed *k*_*max*_ = 40 on the tested GPU devices. This in turn limits the supported database sequence length to 32 · 40 = 1280 per warp (*p* = 32). Although this is sufficient for the majority of protein sequences, there are still significantly longer ones; e.g. the average length of protein sequences in Swiss-Prot is 361 but the longest is 35,213.

In order to overcome this length limitation, we have implemented another specialized CUDA kernel that divides the workload into 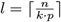 non-overlapping sub-matrices of size *m*×(*k*·*p*) each (except for the rightmost submatrix, which might be smaller) that are computed by a warp in *l* stages from left to right. DP cells in the right column of thread *T*_*p*−1_ are written to CUDA global memory and loaded by thread *T*_0_ in the next stage. In order to avoid repeated, expensive accesses to global memory, these values are only loaded/written every *p* iterations in coalesced fashion. Otherwise, they are exchanged between threads via warp shuffles in a similar way to reading query sequence characters as explained above. Figure 4 illustrates our tiling scheme.

**Figure 4.**
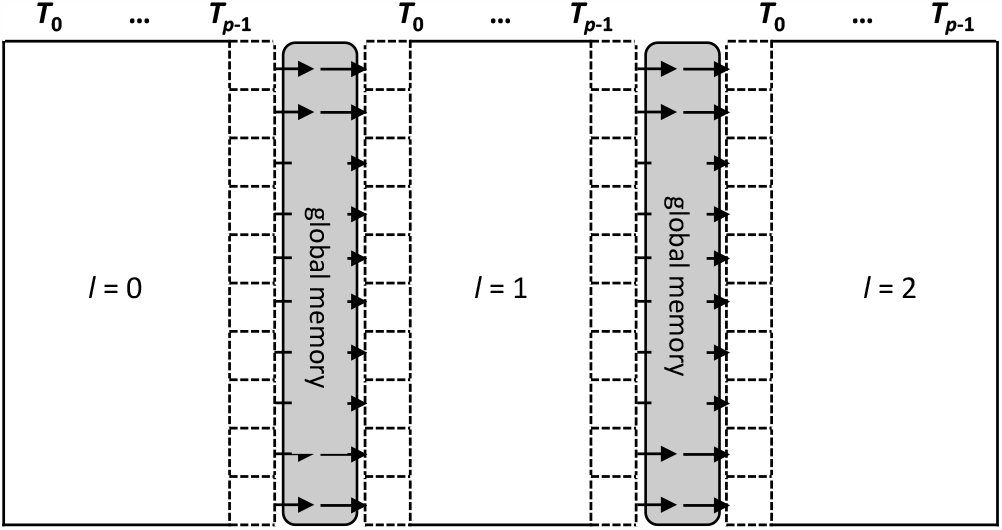
Illustration of our tiling scheme for a database sequence processed in *l* = 3 stages from left-to-right by *p* threads. Each stage compares the full query to one third of the database sequence.

#### Architecture-aware specialized GPU kernels

Using appropriate values of *k* and *p* in our algorithm is critical to optimize execution times. We have thus declared *k* and *p* as C++ template parameters that allows us to generate various GPU kernels tailored to different database sequence lengths ranges and (sub)warp sizes for each GPU architecture generation. For example, using *k* = 32 and *p* = 8 (*k* = 32 and *p* = 4) would generate a kernel that aligns database sequences of length up to 256 (128) with the appropriate number of registers to maximize performance for that combination of sequence length and GPU.

To further evaluate capabilities of various GPU architectures we have implemented score computations using four different (floating-point and integer-based) arithmetic formats:

- `float` (standard 32-bit floating point)
- `half2` (half-precision (16-bit) floating point arithmetic whereby two half-precision values are stored in a 32-bit word)
- `int32` (standard 32-bit integers)
- `s16x2` (two 16-bit integers stored in a 32-bit word)

Ampere, Ada, and Hopper GPUs include micro-architecture improvements and specialized instructions that can be used to accelerate DP algorithms. All described implementations support these features.

Our `half2` and `s16x2` kernels compute two independent alignments at the same time in the lower and the upper part of a packed pair of 16-bit values thus effectively doubling the number of cells computed per thread. Besides instructions for packing (extracting) two 32-bit values into (from) a pair of 16-bit values, we used addition and maximum operations that operate on pairs of 16-bit values to compute the DP recurrence relations.

Kernels are low-level optimized using `Math API` intrinsics to minimize the number of executed instructions in the inner-loop. We have validated that the compiler generates the best combination of instructions for each architecture obtaining high performance.

Our implementation of integer-based kernels can take advantage of DPX instructions which are hardware-accelerated on Hopper GPUs and simulated in software on other GPUs. Specifically, for `s16x2` we use the following DPX instructions

- `__viaddmax_s16x2(a,b,c)` (performs per-halfword max(*a* + *b, c*)), and
- `__vimax3_s16x2 relu(a,b,c)` (performs per-half-word max(max(max(*a, b*), *c*), 0)).

Additionally, our half-precision implementation takes advantage if the `vhmnmx` instruction on Ada GPUs allowing max operations with three half2 input operands using only a single instruction.

Note that rounding issues can occur when using `half2` in cases where scores in the DP matrix exceed 2,048. If this happens, we mark the corresponding database sequence and the alignment will be re-computed using 32-bit precision arithmetic. Note that this re-computation is typically only required for a few sequences of the database and corresponding performance losses are negligible. In a similar way we mark overflowing sequences for kernels with the s16x2 datatype if scores exceed the maximal value for signed 16-bit integers (32,767). This fallback re-computation strategy is also GPU-accelerated.

### Batch Partitioning and Sorting

PCIe data transfers can be performed concurrently to GPU-based alignment computation by partitioning the protein sequence database into batches. Using CUDA streams we overlap the computation of the current batch and the transfer of next batch from host to GPU.

Once the current batch has been transferred to the GPU, it is further partitioned with respect to predefined sequence length ranges. Each partition corresponds to a specific alignment kernel that is optimized for a certain database sequence length range based on using different combinations of the template parameters *p* and *k*. A typical example may feature 20 kernels tailored for the length ranges [64 · *i* + 1, 64 · (*i* + 1)] for 0 ≤ *i* ≤ 19 plus a kernel with DP matrix partitioning for all database sequences exceeding 1280 in length. Each of these kernels is executed concurrently using CUDA streams to align the query to the set of databases sequences within the corresponding partition.

After all batches have been processed, the computed alignment scores are sorted on the GPU. The scores and the ID of the top-ranked database sequences are transferred to the host for output to the user.

### Multi-GPU support

CUDASW++4.0 comes with multi-GPU support. When multiple GPUs are available, the database is split across the GPUs. Each GPU maintains its own result list of the top best scores. In a final processing step, these per-GPU lists are gathered on a single GPU to determine the overall best scores. The associated advantages are two-fold. First, the total available GPU memory increases which allows us to fit larger databases into fast GPU memory which in turn reduces PCIe data transfers. Second, the processing speed per query increases linearly with the number of GPUs since the per-GPU work decreases and multiple GPUs can operate in parallel.

Recall that we use different alignment kernels for different sequence lengths. To reduce work imbalance, we identify the corresponding sequences for each kernel in the database and assign them to the GPUs such that each GPU receives approximately the same number of amino acids in total.

## Results

### Experimental Design

We compared CUDASW++4.0 to two GPU-based protein sequence alignment algorithms (CUDASW++3.0 (v.3.1.1) and ADEPT (v.1.0)) and to three CPU-based algorithms (BLASTP (v.2.9.0+, as part of BLAST+ v.2.14.0), SWIPE (v.2.1.1), and SWIMM2.0 (v.2.1.0)). We excluded SWhybrid and Parasail since they terminated prematurely (SWhybrid) or were too slow (Parasail).

We used 20 protein sequences as queries with lengths ranging from 144 to 5,478 to search against three real-world databases of different size (Swiss-Prot (small), UniRef50 (medium), TrEMBL (large)) and simulated databases of equal-sized sequences (in order to test the achievable peak performance of our kernels):

#### Swiss-Prot

UniProtKB/Swiss-Prot [30] (Rel.: 2023 03) contains 206,858,779 amino acids in 569,793 sequences with largest (average) sequence length of 35,213 (361).

#### UniRef50

UniRef50 [31] (Rel.: 2023 03) consists of 17,340,143,855 amino acids in 60,952,894 sequences with largest (average) sequence length of 45,354 (282).

#### TrEMBL

UniProtKB/TrEMBL [30] (Rel.: 2023 03) contains 86,972,040,633 amino acids in 248,272,897 sequences with largest (average) sequence length of 45,354 (348).

#### Simulated

The simulated databases consist of 1M sequences each of length 128, 256, 512, 1024, and 2048, respectively (named D128, D256, D512, D1024, and D2048).

The accession numbers of all queries are: P02232, P05013, P14942, P07327, P01008, P03435, P42357, P21177, Q38941, P27895, P07756, P04775, P19096, P28167, P0C6B8, P20930, P08519, Q7TMA5, P33450, and Q9UKN1, listed in the ascending order of sequence length.

All SW-based tools are executed using the scoring scheme with BLOSUM62, *α* = 11, *β* = 1. For comparison to BLASTP we also tested performance for two different common scoring schemes with BLO-SUM62, *α* = 11, *β* = 1 (BL62) and with BLOSUM50, *α* = 13, *β* = 2 (BL50). Note that the runtimes for SW-based tools are almost identical for both scoring schemes, but can vary significantly for BLASTP.

Speeds for SW-based tools are reported by measuring runtimes and converting them into the number of DP matrix cell updates that are performed per second; i.e., **TCUPS** (<u>T</u>rillions of <u>C</u>ell <u>U</u>pdates <u>P</u>er <u>S</u>econd).

Experiments were conducted on the following systems:

#### S1 (L40S, Ada)

AMD Ryzen Threadripper 3990X CPU (64 physical cores, 128 logical cores) 2.9 GHz, L40S GPU with 48GB GDDR6 device memory.

#### S2 (A100, Ampere)

Dual-socket AMD EPYC 7713 CPU (64 physical cores, 128 logical cores), 2.9 GHz, 8 A100 GPUs (SXM), each with 80GB HBM2e device memory.

#### S3 (H100, Hopper)

Dual-socket Xeon Gold 6130 CPU (32 physical cores, 64 logical cores), 2.1 GHz, 8 H100 GPUs (SXM) each with 80GB HBM3 device memory (pre-production).

BLASTP and SWIPE are executed on S1 using 128 CPU threads, while SWIMM2.0 is executed on S3 using 64 threads since it supports AVX-512 instructions only available on Intel systems. GPU experiments have been conducted using a single GPU on the systems S1, S2, and S3. Furthermore, multi-GPU scalability tests are performed on S2 and S3 for up to 8 GPUs. nvcc 12.4 (internal pre-production) and gcc 9.3.0 are used as GPU and host compiler, respectively.

### Kernel Peak Performance and efficiency

We first evaluate the performance of our CUDA kernels in an ideal situation where all database sequences are of same length. To identify the maximal achievable performance we just measure kernel execution times; i.e. we exclude any overheads like PCIe data transfers and we do not consider varying database sequence lengths.

Table 1 shows the achieved peak performance in terms of TCUPS for the four different datatypes (float, half2, int, s16x2) on the three tested GPU architectures for scanning the simulated databases with each of the 20 queries. H100 achieves the highest performance with the overall peak of 5.71 TCUPS using DPX s16x2. DPX instructions only have native support on H100 while they are only emulated on A100 and L40S. Thus, H100 is 6.2x and 13.6x faster compared to L40S and A100 for this datatype, respectively. DPX s16x2 also doubles the performance compared to DPX s32 on the same platform as expected.

**Table 1.** Kernel peak performance in TCUPS (Tera Cell Updates per Second) for scanning simulated databases with equal-length protein sequences for different datatypes on three different GPU architectures.

| Datatype | H100 | L40S | A100 |
| --- | --- | --- | --- |
| s16x2 | **5.71\*** | 0.92 | 0.42 |
| int | **2.85\*** | 1.97 | 0.99 |
| half2 | 4.43 | **5.01** | 1.94 |
| float | 2.14 | **2.58** | 1.28 |

The half2 datatype is fastest on both L40S and A100, achieving 5.01 and 1.94 TCUPS, respectively. L40S provides 2.6x higher performance compared to A100, this is mainly due to (a) dual-FP port SM resources, (b) 20% reduction of executed max instructions by using `vhmnmx`, (c) higher clock frequency (1.5x), and (d) higher SM count (1.31x) compared to A100.

Note that the required half2 max comparison function __`hmax2()` is only natively supported on Ampere GPUs and successor generations and thus our half2 kernels would run significantly slower on older GPU generations like Volta.

We now analyze if our approach is able to effectively remove overheads occurred by memory accesses using half2 and s16x2 arithmetic as an example by modeling the *theoretical peak performance* (TPP) in terms of cell updates per second (CUPS) of the utilized GPU hardware as:

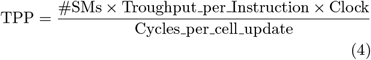

Cycles_per_cell_update in Eq. 4 models the maximum attainable performance constrained by the algorithm structure and the specifications of the architecture [32]. In our case it refers to the minimal number of clock cycles needed by an individual core of the utilized GPU to calculate one DP matrix cell in *H*. The number of required arithmetic operations in our implementation to calculate a single DP cell is 10 (i.e. 6 maximum instructions and 4 additions/subtractions). These values are determined by Eq. 1 (1 add, 3 max), optimized versions of Eq. 2 and 3 (3 sub, 2 max)^[2]^, and a comparison to the running maximum value (1 max). This leads to 10 required clock cycles per cell update (when using single-precision arithmetic) and 5 required clock cycles (when using half2 arithmetic).

Table 2 shows the achieved efficiencies for the best-performing datatype on each GPU. Note that we disregard the lookup operation *σ*(*q*_*i*_, *s*_*j*_) or any value to register movements, since it could in principle be performed concurrently to computation. Additionally, other important low-level considerations as ALU/FP dual-port, max3 and DPX fused operations that allow for the reduction of the number of instructions have been taken into consideration in the efficiency calculation.

**Table 2.** Efficiency of CUDASW++4.0 for best performing kernels per platform. Achieved TCUPS is the best performance in Table 1 (TPP = theoretical peak performance).

|  | H100 | L40S | A100 |
| --- | --- | --- | --- |
| Stream-Multiprocessors (SMs) | 132 | 142 | 108 |
| Max Clock (GHz) | 1.98 | 2.56 | 1.41 |
| Sustained Clock (GHz) | 1.98 | 2.11 | 1.41 |
| TPP (Max Clocks) TCUPS | 8.36 | 7.74 | 1.95 |
| TPP (Sustained Clocks) TCUPS | 8.36 | 6.38 | 1.95 |
| Achieved TCUPS | 5.71 | 5.01 | 1.94 |
| Efficiency (Max Clocks) | 68% | 65% | 99% |
| Efficiency (Sustained Clocks) | 68% | 79% | 99% |
| Power Consumption (real/TDP) | 554/700w | 347/350w | 348/350w |
| Datatype | s16x2 | half2 | half2 |
| Maximum Cell Representation | 16 bits | 11 bits | 11 bits |

After optimizations, the inner loop of the H100 implementation is composed of just 5 max/add instructions plus 2 instructions for data movement to calculate 64 DP cells. This represents a theoretical 71% efficiency for a real implementation limited by just instruction execution and considering the 2 data movements overhead for the actual SW calculation. Effective software stack, compiler and code generation are key to achieve this level of efficiency. Any additional or mis-scheduled instructions will have significant impact on the final performance.

Our approach is able to achieve efficiencies up to 68% on H100 and 99% on A100 GPU architectures, which shows that the warp-shuffle tiled approach is able to effectively transform the problem to compute-bound and minimize overheads from memory accesses.

### Kernel Performance for Varying Sequence Lengths

Figure 5 shows the peak performance of our kernels in terms of TCUPS for scanning the simulated databases (D128, D256, D512, D1024, D2048) on H100 (s16x2), L40S (half2), and A100 (half2) with each of the 20 queries. Database sequences with lengths up to 1024 were aligned without the need of tiling the DP matrix while our tiling scheme is used for D2048. Performance for D2048 only decreases by around 7.0%, 1.9%, and 5.5% compared to D1024 on H100, L40S and A100, respectively, due to additional writing and reading to and from global memory. This shows that our generalization of the DP matrix tiling scheme for arbitrary sequence size is scalable and efficient. For the kernels without matrix tiling, we noticed that maximal performance is achieved when computing *k* = 32 values per thread and varying the sub-warp sizes according to the DB sequence length (i.e. we use *p* = 4, 8, 16, 32 for D128, D256, D512, D1024, respectively). Increasing the number of values per thread beyond 32, however, generally decreases performance due to register pressure.

**Figure 5.**
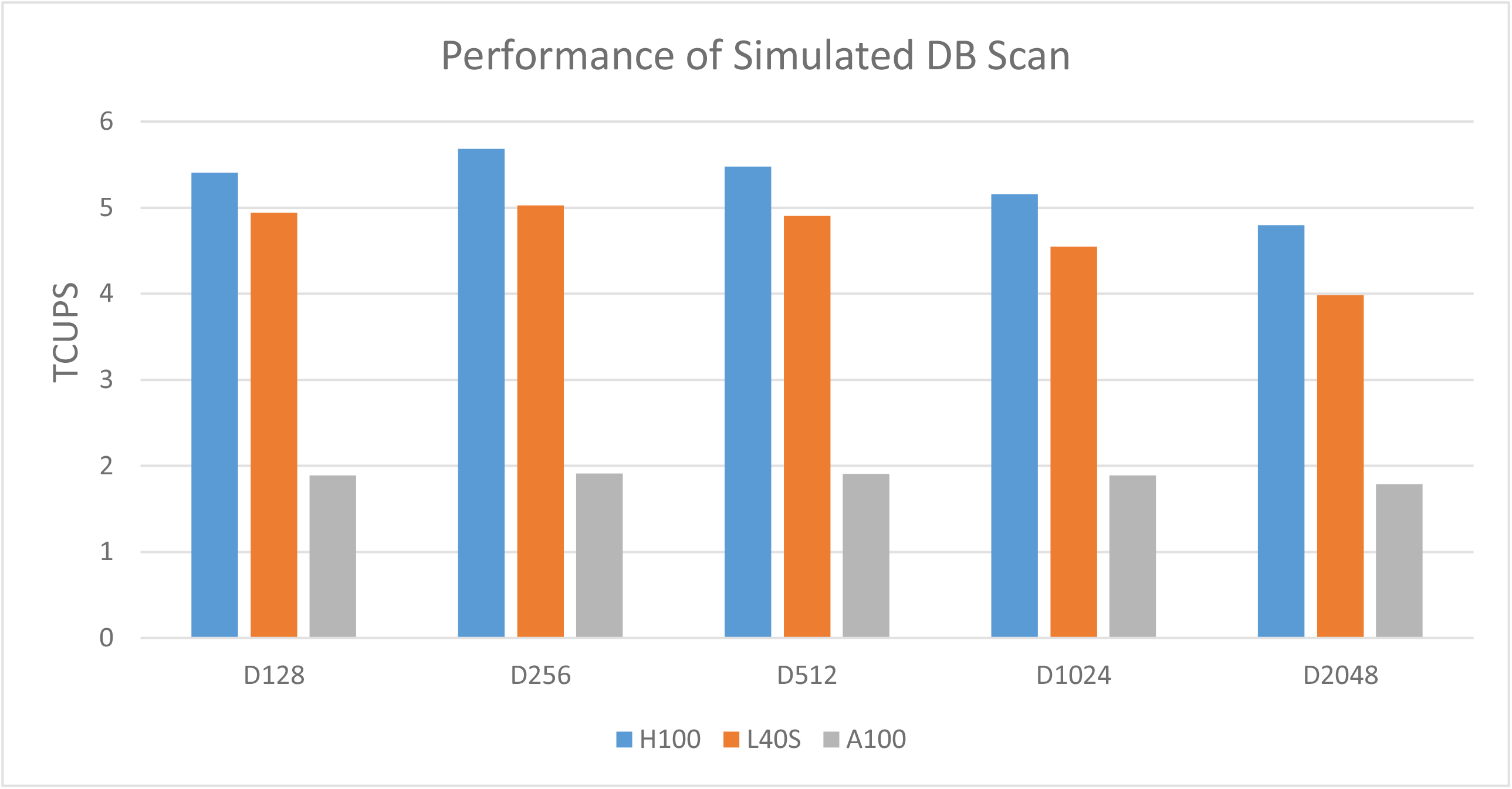
Peak performance (in terms of TCUPS) of our alignment kernels for scanning databases with subject sequences of identical length (D128, D256, D512, D1024, D2048) on H100 (using s16x2 kernels), L40S (using half2 kernels), and A100 (using half2 kernels).

Figure 6 shows the performance in terms of TCUPS for scanning Swiss-Prot, UniRef50, and TrEMBL with the set of 20 queries of various length using s16x2 kernels on H100. This tests the performance for real-word scenarios with a typical distribution of sequences of different lengths. We execute several kernels tailored for different length ranges; i.e separate kernels for all subject sequences with the length range [64·*i*+1, 64·(*i*+1)] for *i* = 0, …, 19 using corresponding values of *k* and *p* are executed. In addition kernels designed for longer sequences (i.e. with DP matrix tiling) are executed for all subject sequence exceeding 1280 amino acids. Measured GPU runtimes now also include PCIe data transfer times in addition to various other overheads (tiling, partitioning, sorting, recomputation). Note that all database sequences can be stored in GPU device memory for Swiss-Prot and UniRef50. Thus, they are only transferred once for all 20 scan tasks, which leads to lower performance for the first query. For TrEMBL the whole database does not completely fit into GPU memory and is thus transferred for each query. Thus, the performance of shorter queries is again lower due to the additional PCIe data transfer overhead. However, for all queries of at least 375 in length performance consistently exceeds 4.4 TCUPS which shows that PCIe data transfers can be effectively overlapped with computation. The results also indicate that kernel performance generally increases for longer queries since the relative overhead at the beginning and the end of the wavefront decreases.

**Figure 6.**
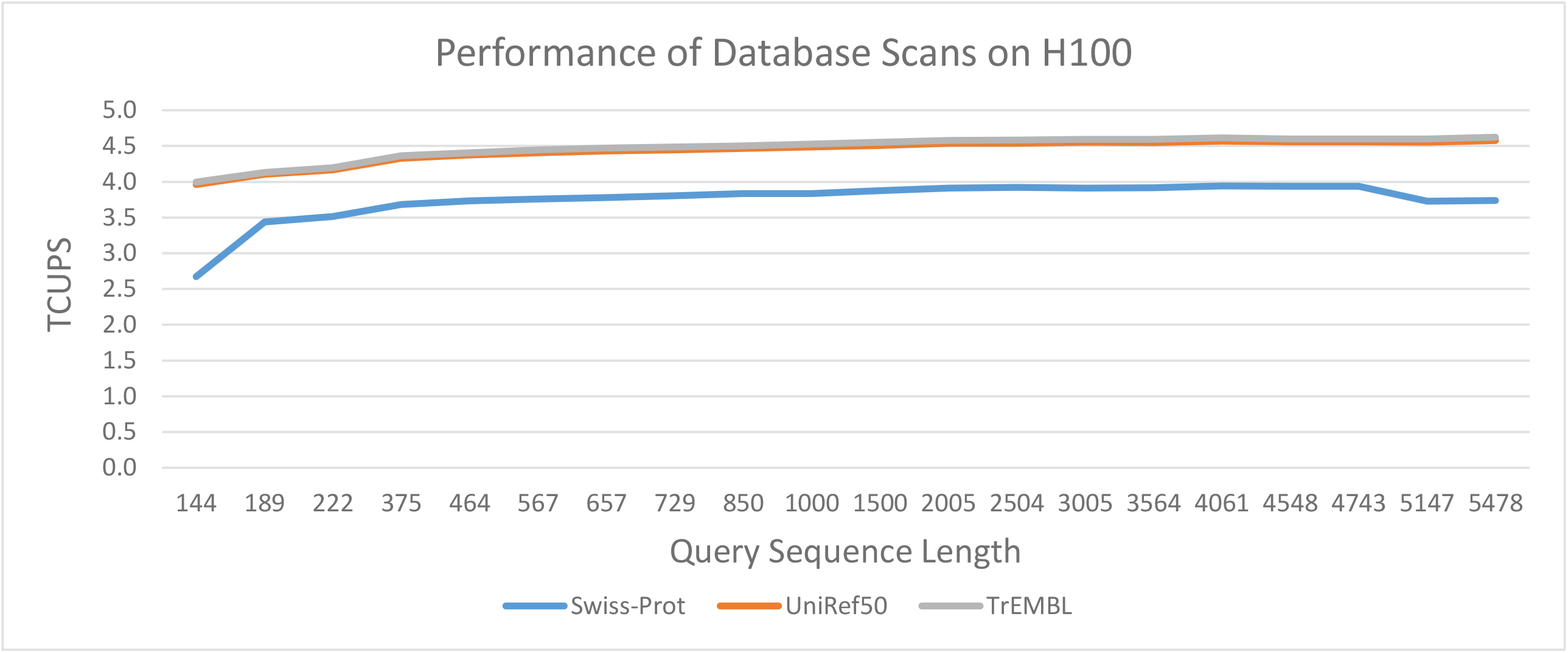
Performance (in terms of TCUPS) for scanning Swiss-Prot, UniRef50, and TrEMBL with queries of different length on H100.

Despite highly varying sequence lengths in the databases and the various other overheads relatively stable performance of up to 4.0 TCUPS, 4.6 TCUPS, and 4.7 TCUPS can be achieved for scanning Swiss-Prot, UniRef50, and TrEMBL on the H100, respectively. This corresponds to 70%, 81%, and 83% of the overall peak kernel performance for equal subject sequence lengths without any overheads.

### Performance Comparison to other tools

Figure 7 compares the performance of CUDASW++4.0 to other SW-based tools on GPUs (CUDASW++3.0, ADEPT), and CPUs (SWIPE, SWIMM2.0) for scanning UniRef50 with queries of different lengths in terms of TCUPS. Measured GPU runtimes include PCIe data transfer times in addition to various other overheads (partitioning, sorting, recomputation). SWIPE is executed using 128 threads on the AMD Ryzen Threadripper 3990X CPU, while SWIMM2.0 is executed on a 32-core Intel Xeon Gold 6130 CPU using 64 threads to evaluate the impact of using AVX-512 instructions. All GPU-based tools were executed on an H100. Note that ADEPT was not able to process queries longer than 1,000. All tools use the same scoring scheme (BLOSUM62, *α* = 11, *β* = 1).

**Figure 7.**
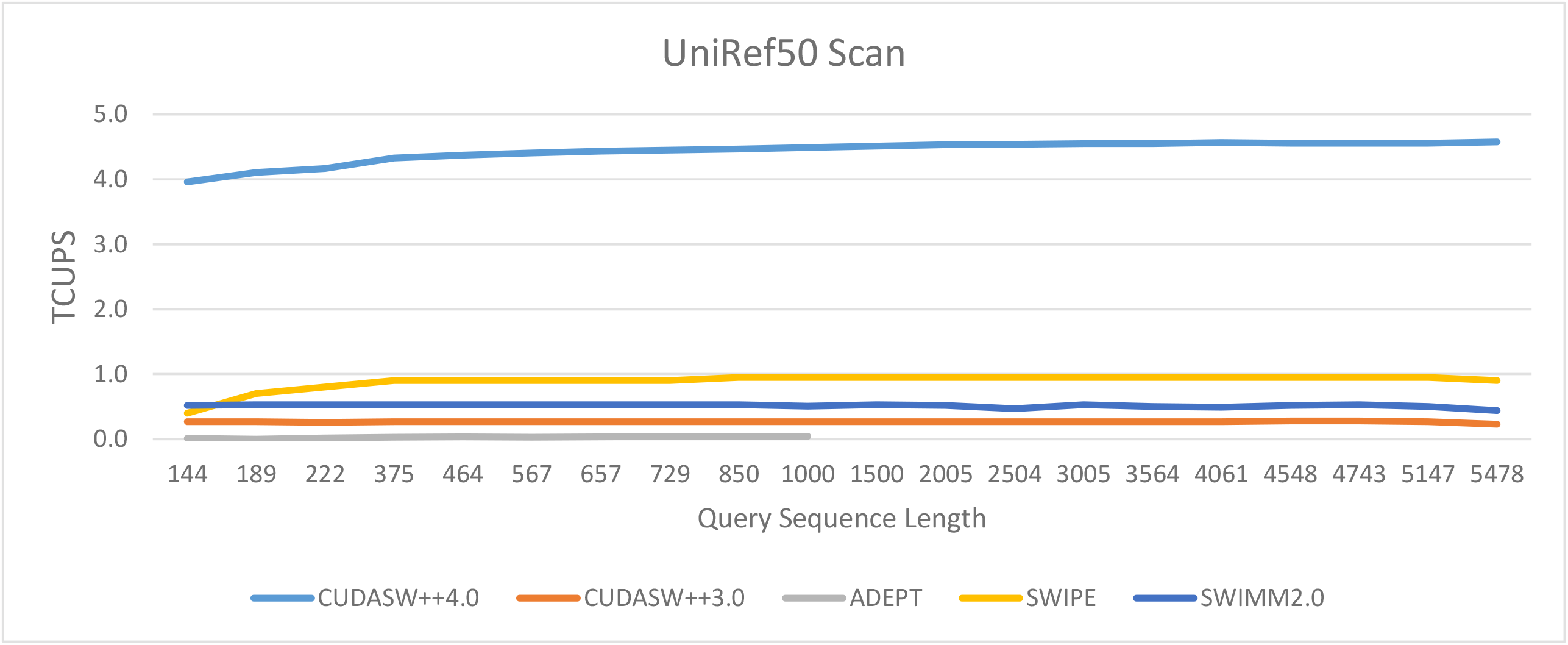
Performance (in terms of TCUPS) for scanning the UniRef50 database with queries of different length using different SW-based tools on GPUs and CPUs.

CUDASW++4.0 yields an average performance of 4.57 GCUPS and is superior to all other tested SW-based tools for every query. It runs on average 16.2×, 134x, 4.9x, and 8.5x faster than CUDASW++3.0, ADEPT, SWIPE, and SWIMM2.0, respectively.

Different from SW-based tools, the performance of BLASTP can strongly fluctuate for different scoring schemes and different query lengths. Thus, we measure the full execution times of BLASTP for the default settings for BLOSUM62 (*α* = 11, *β* = 1) and BLOSUM50 (*α* = 13, *β* = 2) for all 20 query using 128 threads on the AMD Ryzen Threadripper 3990X CPU using the TrEMBL, UniRef50, and SwissPort databases. Those are compared to the full execution times of CUDASW++4.0 on H100 using again both scoring schemes. The results are shown in Table 3. Overall, CUDASW++ outperforms BLASTP for scanning TrEMBL, UniRef50, and Swiss-Prot by a factor of 1.7, 1.9, and 1.7 for the BL62 scoring scheme and by a factor of 7.5, 7.9, and 5.5 for the BL50 scoring scheme, respectively.

**Table 3.** Full execution times and corresponding speedups for scanning TrEMBL, UniRef50, and Swiss-Prot with 20 query sequences of CUDASW++4.0 on H100 and BLASTP 2.0.9+ (from BLAST+ 2.14.0) on AMD Ryzen Threadripper 3990X using 128 CPU threads using the BLOSUM62 scoring scheme (BL62) and the BLOSUM50 scoring scheme (BL50).

|  |  | TrEMBL |  | UniRef50 |  | Swiss-Prot |  |
| --- | --- | --- | --- | --- | --- | --- | --- |
|  |  | BLASTP | CUDASW4 | BLASTP | CUDASW4 | BLASTP | CUDASW4 |
| BL62 | Runtime | 22m 27s | 13m 9s | 4m 58s | 2m 37s | 5.3s | 2.9s |
|  | Speedup |  | 1.7x |  | 1.9x |  | 1.8x |
| BL50 | Runtime | 100m 50s | 13m 6s | 21m 26s | 2m 36s | 17s | 2.9s |
|  | Speedup |  | 7.7x |  | 8.2x |  | 5.9x |

### Multi-GPU Scaling

To evaluate scalability with respect to the number of used GPUs connected to the same host, we measured the total execution times of scanning the TrEMBL database for our 20 queries on Systems S2 and S3, using 1, 2, 4, and 8 GPUs, respectively. With only a single GPU, the database does not fit into GPU memory and needs to be processed in batches which are transferred over PCIe for each query sequence. When two or more GPUs are used, the database fits into GPU memory and we only need to transfer it once for all queries.

Table 4 presents our results in terms of runtime, speedup, and parallel efficiency for A100 and H100 multi-GPU systems. CUDASW++4.0 shows good scaling with respect to the number of GPUs, achieving a speedup of 7.7 with 8 H100 GPUs which corresponds to an efficiency of more than 90%.

**Table 4.** Full execution times, speedups, and parallel efficiency for scanning TrEMBL with 20 query sequences of CUDASW++4.0 using multiple A100 and H100 GPUs.

| Number of GPUs | 1 | 2 | 4 | 8 |
| --- | --- | --- | --- | --- |
| A100 Runtime | 34m 12s | 17m 30s | 8m 57s | 4m 40s |
| A100 Speedup | 1 | 1.95 | 3.82 | 7.33 |
| A100 Parallel Efficiency | 100.0% | 97.7% | 95.5% | 91.6% |
| H100 Runtime | 15m 5s | 7m 33s | 3m 49s | 1m 57s |
| H100 Speedup | 1 | 1.99 | 3.95 | 7.73 |
| H100 Parallel Efficiency | 100.0% | 99.7% | 98.6% | 96.5% |

The slight drop in parallel efficiency for larger number of GPUs can be explained by a certain degree of load imbalance. Recall that our databases are partitioned by sequence lengths, where each partition is distributed amongst GPUs. Some GPUs may thus end up with longer sequences than others which can lead to load imbalance and some loss of parallel efficiency for that partition. Nevertheless, our evaluation shows that our approach is still efficient for up to 8 GPUs (e.g. 91.6% and 96.5% parallel efficiency for 8xA100 and 8xH100, respectively). However, for even larger systems (such as GPU clusters) more sophisticated schemes might be required.

### Energy efficiency

We conducted an energy efficiency analysis on H100 and L40S GPUs. Our experiments use the `nvidia-smi` power governor to limit the maximum power consumption that GPUs can operate. Notice that this setup does not require any code changes, it relies on the driver and is fully transparent to the user.

Fig. 8 shows the performance in TCUPS on H100 and L40S for the maximum operational power consumption ranging from 100 to 700 Watts for scanning a synthetic database with equal-sized sequences. L40S can deliver 0.85 TCUPS running at only 100 Watts. This is similar to standard FPGA energy consumption range, but at higher performance. H100 never consumes more than 550 Watts which is under its 700-Watt TDP. This provides high-performance alignment processing at a low budget energy consumption. The best energy efficiency is delivered by H100 at 15.7 GCUPS/Watt as shown in Fig. 9. This is 23x higher than OSWALD [17] – the most recent FPGA-based implementation for SW protein database scanning [34].

**Figure 8.**
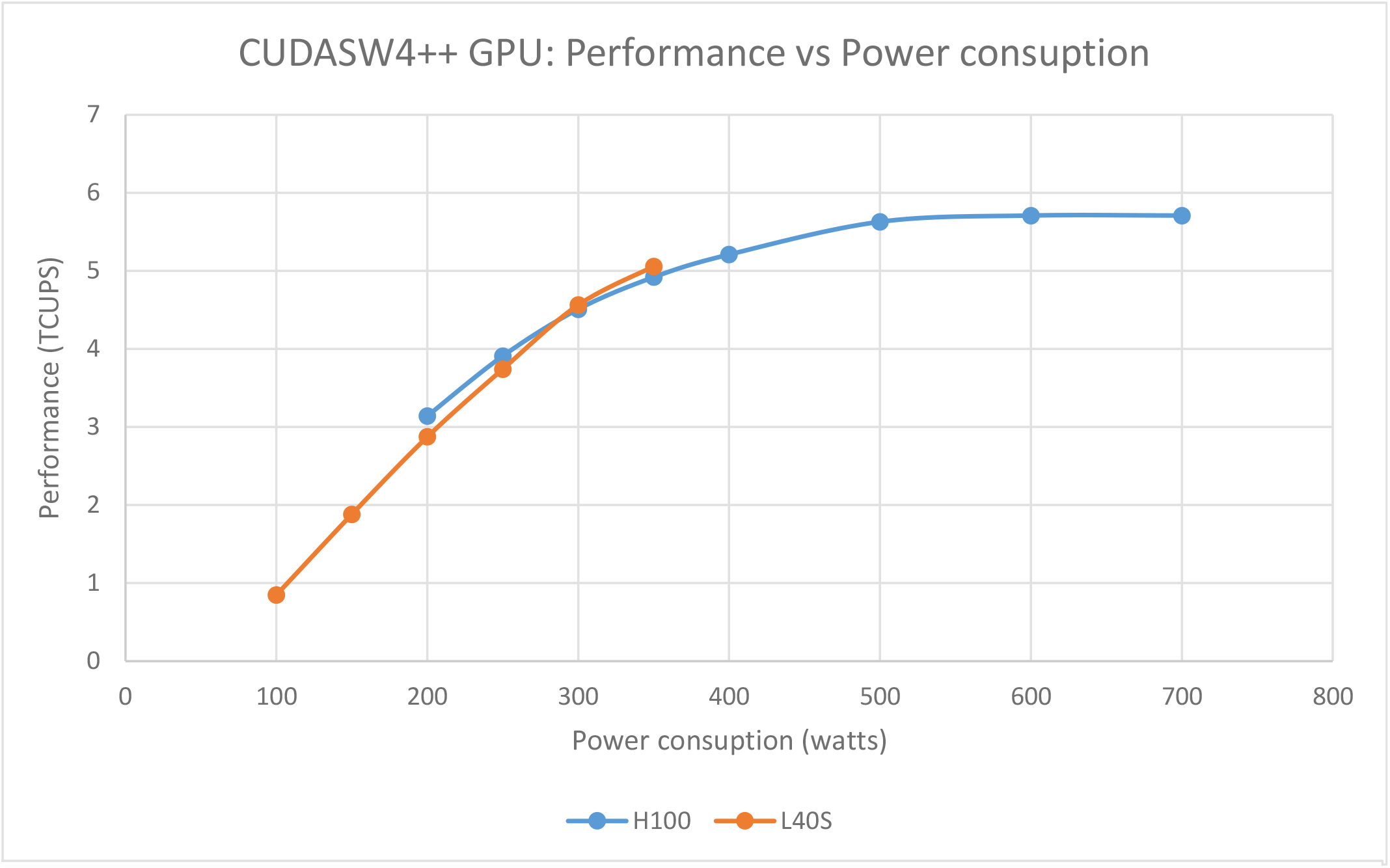
Peak Performance (TCUPS) of alignment kernels scanning synthetic databases with all sequences identical size setting an specific power consumption limit using L40S and H100.

**Figure 9.**
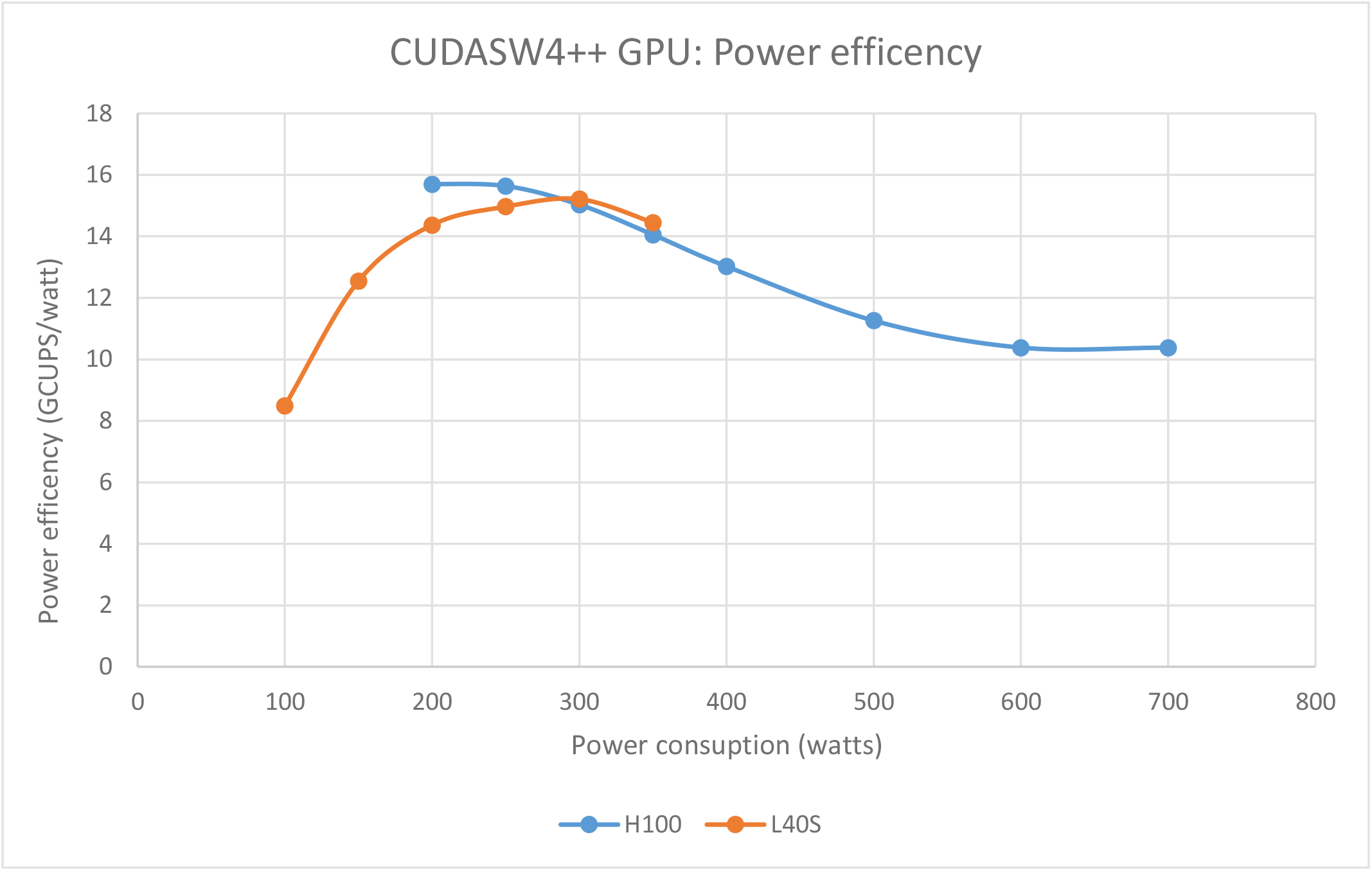
Performance normalized by power consumption running synthetic databases on H100 and L40S (all sequences same size).

## Conclusions

The continually increasing volume of sequence data has resulted in a growing demand for fast implementations of core algorithms. Computation of optimal local pairwise alignments based on DP is an important algorithmic part of many protein sequence analysis tasks and a major contributor to overall runtime due to its quadratic time complexity.

In this paper, we have presented CUDASW++ 4.0, a fast SW-based protein database search software tool, which gains high speed on modern GPUs architectures based on using warp shuffles for communicating data between threads and exploiting the latest GPU features.

Our implementation achieves over 65% of the theoretical peak performance for kernels with s16x2 and half2 arithmetic on three tested GPUs of different generations (Ampere, Hopper, Ada) and outperforms state-of-the-art GPU-based competitors such as CUDASW++3.0 and ADEPT by over an order-of-magnitude.

Furthermore, CUDASW++4.0 offers a clear advantage over state-of-the-art CPU-based SW approaches. Using a single H100 it outperforms SWIPE and SWIMM2.0 running on a 64-core AMD workstation with 128 threads and 32-core Intel workstation with 64 threads by 4.9× and 8.5× on average, respectively. Moreover, our comparison to BLASTP reveals speedups between 1.7 and 1.9 for the BL62 scoring scheme and between 5.5 and 7.9 for the BL50 scoring scheme. Furthermore, our approach can exploit multiple GPUs connected to the same host in order to provide even higher speeds with close-to-linear scaling with respect to the number GPUs.

Our results show that prior GPU approaches like CUDASW++3.0 and ADEPT have not been able to unlock the full potential of GPUs for protein sequence alignment and only achieve a fraction of the available peak performance. As a consequence, they cannot provide significant speedups compared to current CPU-based approaches since they are bottlenecked by inefficient memory access schemes. Our approach thus changes the standing of GPUs in pairwise protein sequence alignment significantly. We demonstrate that the SW method can be re-designed to be fully compute bound, reaching close-to-peak performance on modern GPUs architectures, with a peak performance of over 5.7 TCUPS on an H100. Our introduced schemes for DP matrix tiling and sequence batch partitioning allow for efficient alignment of a variety of sequence lengths that occur in typical protein sequence databases. Thus, GPUs can be efficiently used for typical problem sizes occurring in common databases like Swiss-Prot, UniRef, and TrEMBL.

We have further evaluated different techniques to reduce the amount of executed instructions and the impact of using different datatypes in the DP algorithm (float, half2, int32, and s16x2). Our results show that integer-based datatypes (specifically s16x2) can achieve higher performance on an H100 when taking advantage of the recently introduced DPX instructions. Additionally, we have evaluated the power consumption of CUDASW4++ on H100 and L40S GPUs, achieving up to 15.7 GCUPS/Watt, and demonstrated that GPUs can be more an order-of-magnitude more energy efficient than FPGAs and under the same maximum power budget.

Our parallelization scheme may also be applicable for accelerating iterative profile searching and clustering of large-scale protein sequence sets [35] on GPUs and could be extended to a wider range of DP-based algorithms with similar dependency relationships such as the Viterbi algorithm for profile hidden Markov models [36] or computation of multiple sequence alignments based on pairwise alignments [37]. It would thus be interesting to evaluate the performance of our approach when adapted to different DP algorithms.

The performance and energy efficiency levels presented in this paper will make a direct impact to the future design of data center environments, and as well, on-instrument medical devices for genomic analysis, identifying GPUs a solid alternative to current CPU and FPGA platforms for radically changing the scenarios with these order-of-magnitude cost-, energy-, and time-reduction factors.

## Availability and requirements

Project name: CUDASW++4.0

Project home page: https://github.com/asbschmidt/CUDASW4

Operating system(s): Linux

Programming language: C++, CUDA

Other requirements: CUDA toolkit ≥ 12

License: Apache-2.0

## List of abbreviations

GPU: Graphics Processing Unit
SW: Smith-Waterman
DP: Dynamic Programming
BLAST: Basic Local Alignment Search Tool
SM: Streaming Multiprocessor
SIMT: Single Instruction Multiple Threads
DB: Database
FP: Floating Point
TCUPS: Tera Cell Updates Per Second
FPGA: Field Programmable Gate Array

## Ethics approval and consent to participate

Not applicable.

## Consent for publication

Not applicable.

## Acknowledgements

Not applicable

## Availability of data and materials

Real-world datasets are publicly available.

- Swiss-Prot: https://ftp.expasy.org/databases/uniprot/current_release/knowledgebase/complete/uniprot_sprot.fasta.gz
- UniRef50: https://ftp.expasy.org/databases/uniprot/current_release/uniref/uniref50/uniref50.fasta.gz
- TrEMBL: https://ftp.expasy.org/databases/uniprot/current_release/knowledgebase/complete/uniprot_trembl.fasta.gz
- Query protein sequences: https://github.com/asbschmidt/CUDASW4/allqueries.fasta

## Authors’ contributions

BS and FK are the main developers of CUDASW++4.0. BS, FK, CH and AC conducted the experiments and optimizations. BS proposed and supervised the project. BS, FK, CH, and AC contributed to the manuscript. All authors read and approved the final manuscript.

## Footnotes

[1] On our systems parsing time was reduced from around 30 seconds for UniRef50 in FASTA to only 0.3 seconds for reading the customized local database.

[2] Subtractions in Eqs. 2 and 3 can be reduced from 4 to 3 since _*β*_ (or _*α*_) only needs to be subtracted once (using _max*{E*(*i*−1, *j*), *F* (*i, j* −1)*}*−*β*_) and keeping a temporal _*α*+*β*_ register that is finally subtracted to _*H*_ maximum [3, 33]

